# TIMSCONVERT: A workflow to convert trapped ion mobility data to open data formats

**DOI:** 10.1101/2021.12.09.472024

**Authors:** Gordon T. Luu, Itzel Lizama-Chamu, Catherine S. McCaughey, Laura M. Sanchez, Mingxun Wang

## Abstract

**Summary:** Advances in mass spectrometry instrumentation have led to the development of mass spectrometers with ion mobility separation (IMS) capabilities and dual source instrumentation, but the current software ecosystem lacks interoperability with downstream data analysis using open-source software/pipelines. Here, we present TIMSCONVERT, a data conversion workflow from timsTOF fleX MS raw data files to size conscious mzML and imzML formats with minimal preprocessing to allow for compatibility with downstream data analysis tools, which we showcase with several examples using data acquired across different experiments and acquisition modalities on the timsTOF fleX.

**Availability and Implementation:** TIMSCONVERT and its documentation can be found at https://github.com/gtluu/timsconvert and is available as a standalone command line interface, Nextflow workflow, and online in the Global Natural Products Social (GNPS) platform (https://proteomics2.ucsd.edu/ProteoSAFe/index.jsp?params={%22workflow%22%3A%20%22TIMSCONVERT%22}).

**Contact:** Mingxun Wang

**Supplementary Information:** Supplementary data are available at *Bioinformatics* online.

## Introduction

In recent years, ion mobility spectrometry (IMS) has been integrated into instruments configured for liquid chromatography-tandem mass spectrometry (LC-MS/MS). Examples include the timsTOF Pro (Bruker Daltonics) featuring an electrospray ionization (ESI) source and timsTOF fleX, which is a dual source instrument configured with ESI and matrix-assisted laser desorption/ionization (MALDI) sources coupled to a trapped ion mobility spectrometer (TIMS) and hybrid quadrupole-time-of-flight (qTOF) mass analyzer. Ion mobility - mass spectrometry (IM-MS) allows for the separation of ions based on their mobility in a carrier buffer gas; a common application includes the separation of isobars (compounds with the same nominal mass but different chemical formula) and isomers (compounds with the same chemical formula but different 3D configuration) in biological samples. The inclusion of IM-MS online separation increases the dimensionality of the data produced during acquisition. Therefore, the introduction of more advanced instrumentation has also been accompanied by novel data formats allowing for more efficient storage of large datasets.

Here, we have developed TIMSCONVERT with three goals in mind: 1) to incorporate ion mobility data into the commonly used mzML format, 2) convert TSF files to mzML or imzML, and 3) convert TDF files to mzML or imzML (Deutsch, 2008, 2010; Martens *et al*., 2011; Römpp *et* al., 2011; Schramm *et al*., 2012). The dual source capabilities of the timsTOF fleX (Bruker Daltonics) allows for a wide range of experiments to be performed which can be broadly categorized into the following: 1) LC-MS/MS, 2) LC-TIMS-MS/MS, 3) MALDI-qTOF dried droplet (DD), 4) MALDI-qTOF mass spectrometry imaging (MSI), 5) MALDI-TIMS-qTOF DD, and 6) MALDI-TIMS-qTOF MSI. A consequence of the versatility of the timsTOF fleX is the amount of different data formats that result from these different acquisition methods (Supplementary Information). At the time of writing, data conversion for LC-TIMS-MS/MS is supported by DataAnalysis (Bruker Daltonics) and Proteowizard MSConvert, but currently available software does not incorporate ion mobility data into the commonly used mzML and mzXML formats without resulting in unwieldy data files (over eighty times increase in size) (Chambers *et al*., 2012). Packages such as AlphaTims, OpenTIMS, TimsPy, and TimsR have been developed to access and convert the data programmatically, but output formats are currently limited to MGF and HDF5 (Willems *et al*., 2021; Łącki *et al*., 2021). Additionally, MALDI-qTOF and MALDI-TIMS-qTOF DD data are currently only supported by DataAnalysis (Bruker Daltonics). Furthermore, export of MSI data to the open source imzML format is currently only possible with the proprietary program SCiLS Lab (Bruker Daltonics). Therefore, TIMSCONVERT bridges proprietary data formats with existing open source data analysis software to afford flexibility and choice in workflows.

## Implementation

### Conversion of LC-TIMS-MS/MS Runs

Functionality for conversion of LC-TIMS-MS/MS to the widely used mzML format in data-dependent acquisition parallel accumulation-serial fragmentation (ddaPASEF) experiments is available (Meier *et al*., 2018). Importantly, we have leveraged the mzML schema to include ion mobility information in the resulting data. We have developed a workflow leveraging the AlphaTims package over a single run or folder containing multiple runs (Willems *et al*., 2021). This workflow provides options to include MS^1^ spectra in the output, encoding notation, and the method of grouping for MS^1^ spectra. Due to the inclusion of ion mobility in LC-TIMS-MS/MS data, the grouping method is important to consider when exporting data as it has arguably the most impact on the resulting data. When grouped by frame/retention time, a single MS^1^ spectrum is exported without modification, with the exception of the addition of ion mobility data in the form of a binary array. Each frame also contains N number of scans/mobility values (*1/K*_*0*_). When grouped by scan/mobility, a single MS^1^ spectrum is split by scan into N numbers of MS^1^ spectra. Willem *et al*. have provided a more in-depth discussion on the dimensionality and structure of the data (Willems *et al*., 2021). MS/MS spectra are exported without modification; mobility values for the selected precursor ions are included. All spectra are written to mzML files using the psims API (Klein and Zaia, 2019).

### Conversion of MALDI-qTOF DD and MSI Runs

In addition to LC-TIMS-MS/MS conversion, TIMSCONVERT also supports conversion of MALDI-qTOF DD and MSI data from TSF to mzML and imzML, respectively. Here, data is loaded using in-house code based on the TIMS-SDK (Bruker Daltonics). For DD data, users can specify the output directory, filename (single file only), encoding notation, and decide whether spectra from different spots should be combined into a single file or exported to individual files. If choosing the latter, a .csv plate map containing sample name(s) is required. Spectra are written using the psims API (Klein and Zaia, 2019).

Conversion of the MSI data provides the option to specify the output directory, filename (single file only), encoding notation, and whether the .imzML should be written in processed or continuous mode. Importantly, when exporting data to “continuous” imzML files, only the first *m/z* array is used. Data is written using the pyimzML package (alexandrovteam).

### Conversion of MALDI-TIMS-qTOF DD and MSI Runs

MALDI-TIMS-qTOF DD and MSI processing is similar to that of non-TIMS data with one exception. DD spectra can be grouped by frame or scan as described for LC-TIMS-MS/MS data. MSI data exported to imzML omits TIMS data; future iterations will incorporate ion mobility data into imzML files.

## Availability

### TIMSCONVERT Availability and Usage

TIMSCONVERT was developed in Python 3.8 and consists of two workflows for data conversion: an LC-MS workflow and a MALDI workflow. The LC-MS workflow is available online at https://proteomics2.ucsd.edu/ProteoSAFe/index.jsp?params={%22workflow%22%3A%20%22TIMSCONVERT%22}, as a standalone workflow via Nextflow, or Python module to be used via the command line interface (CLI) (Di Tommaso *et al*., 2017). The MALDI workflow is currently only available via the CLI; support in the online and Nextflow workflows is currently in development. In its current iteration, TIMSCONVERT only exports centroided spectra; the option to export profile spectra is planned for future releases. The experiment type and acquisition method (ESI or MALDI) are user specified in the CLI workflow.

### Github Source

All source code and documentation can be found at https://github.com/gtluu/timsconvert.

## Conclusion

In this manuscript, we present TIMSCONVERT: a workflow to convert a wide array of data formats generated by the timsTOF Pro, timsTOF SCP and timsTOF fleX instruments (Bruker Daltonics); TIMSCONVERT is also capable of incorporating ion mobility data into mzML format, with incorporation into imzML planned for the future. We have also provided several examples of use cases from which the resulting data housed in open source formats can be used, including but not limited to Global Natural Products Social molecular networking (GNPS) (Wang *et al*., 2016), IDBac (Clark *et al*., 2018, 2019), Cardinal MSI (Bemis *et al*., 2015), and more (Supplemental Information). With the growing usage of ion mobility and the capability of existing open source data formats to house it, we hope to see more development of community standards for ion mobility data representation; we also hope data analysis software and pipelines implement these standards in the future.

## Acknowledgements

This publication was supported by the National Institute of General Medical Sciences Award Number R01GM125943 (LMS), the National Cancer Institute Award Number R01CA240423 (LMS) of the National Institutes of Health, by the National Science Foundation grant 2128044 (LMS), and UC Santa Cruz Startup funds (LMS).

