## Supplemental Information for "TIMSCONVERT: A workflow to convert trapped ion mobility data to open data formats"

*Bruker timsTOF Data Formats*

All Bruker data is housed in a directory with the .d extension. Within the directory, the type of experiment conducted determines the data format. LC-MS/MS experiments use the BAF file format, which is widely supported by both proprietary (i.e. Bruker DataAnalysis, CompassXport) and open source (i.e. Proteowizard MSConvert) data conversion/analysis software (Chambers *et al.*, 2012). Experiments designed to acquire TIMS data (LC-TIMS-MS/MS, MALDI-TIMS-qTOF DD and MSI) use TDF and TDF_BIN file formats, which contain metadata and compressed raw spectral data, respectively. Compared to BAF files, TDF files have the advantage of storing data in a much more efficient manner. Similarly, the TSF and TSF_BIN file formats have recently been developed for MALDI-qTOF DD and MSI experiments and are similar to TDF and TDF_BIN; the main difference is the lack of TIMS data in TSF and TSF_BIN.

*Use Case 1: Molecular Networking for Dereplication of Small Molecules*

*Methods*

*Penicillium atramentosum* RS17 was grown on plate count agar milk salt (PCAMS; 1 g/L whole milk powder, 1 g/L dextrose, 2.5 g/L yeast extract, 5 g/L tryptone, 10 g/L sodium chloride, 15 g/L agar) at room temperature. Spore stocks were harvested at seven days and normalized to an optical density of 0.1 at $600_{nm}$ ($OD_{600}$) in 1X phosphate buffer solution (PBS) and stored at -80 °C for subsequent experiments.

*P. atramentosum* RS17 working stock (200 µL) were spread onto 200 mL of PCAMS medium in a Nalgene (14.25″ x 12.25″ x 5″) polypropylene tray and cultured at room temperature (RT) for seven days. On day 7, agar was removed from the tray and placed into a 2 L culture flask with 1 L of deionized water and sonicated for 30 minutes. Agar was subsequently removed via vacuum filtration and 20 g of Amberlite XAD16N resin was added to the liquid culture and shaken for 1 h at 225 rpm at RT. Resin and cell mass were collected via vacuum filtration and back extracted with 50:50 dichloromethane:methanol by shaking under the same conditions. The filtrate was dried *in vacuo.* Crude biomass was fractionated via a $C_{18}$ 5g/20mL solid-phase extraction (SPE) cartridge (Supelco) with a step gradient of 40 mL MeOH/Milli-Q $H_2O$ mixtures (10% MeOH, 20% MeOH, 40% MeOH, 60% MeOH, 80% MeOH, 100% MeOH) and 100% ethyl acetate (EtOAc). Henceforth, these fractions will be referred to as RS17A-G. Fractions were dried *in vacuo* and stored at -8 °C.

High resolution LC-TIMS-MS/MS data were collected on a Bruker timsTOF fleX with TIMS separation enabled in positive mode. The detection window was set from 100 to 2000 Da and the TIMS range ($1/K_0$) was set from 0.45 to 1.45. An Agilent Poroshell 120 Å 2.1 x 50mm UPLC column was used with a flow rate of 0.5 mL/min where solvent A was MilliQ $H_2O$ w/ 0.1% formic acid and solvent B was ACN w/ 0.1% formic acid. Data were acquired in Compass Hystar 5.1 and otofControl 6.2. The following gradient was used: hold 5% B from 0 to 2 min, 5-100% B from 2 to 12 min, and hold 100% B from 12 to 14 min, followed by a 1 min equilibration time. For each sample, 2 µL of a 1 mg/mL solution was injected. The ESI conditions were set with the capillary voltage at 4.5 kV. For MS/MS, dynamic exclusion was used and the top nine precursor ions from each $MS^1$ scan were subjected to collision energies scaled according to mass, mobility, and charge state for a total of nine data dependent $MS^2$ events per $MS^1$ via ddaPASEF.

*Results and Discussion*

The inclusion of TIMS in LC-TIMS-MS/MS instruments allows for an online orthogonal method of separation in addition to LC. In addition to separation, TIMS provides insights into the structural properties of small molecules being analyzed. One application in which structural information proves useful is dereplication, or the process of identifying known unknowns, using

Global Natural Products Social (GNPS) molecular networking, which is performed by matching sample fragmentation (MS/MS) spectra to those found in GNPS spectral libraries. Here, RS17E and RS17F were uploaded to GNPS and the classical molecular networking workflow was performed to allow for small molecule annotation and dereplication. The resulting job can be found at https://gnps.ucsd.edu/ProteoSAFe/status.jsp?task=941a77d560f7401dbd8c94c83c65df1c. This led to the detection of meleagrin (*m/z* 434.182 $[M+H]^+$). The molecular network generated from this dataset also detected four structurally related analogues of meleagrin (**Figure S1**); based on the mass differences, the following predictions were made; the analogue *m/z* 464.187 was the result of an addition of 30.005 Da, indicating a possible occurrence of meleagrin O-methylation. We hypothesize *m/z* 448.198 (+14.016 Da, addition of methylene group) to be oxaline which is further supported by the study performed by Newmister *et al*. in which oxaline was detected from the marine-derived *Penicillium oxalicum* F30 (Newmister *et al.*, 2016). Similarly, *m/z* 470.178 appears to be the sodiated adduct ($[M+Na]^+$) of oxaline. Lastly, *m/z* 478.171 appears to be an unknown analogue with an addition of 29.973 Da, which we believe to be O-methylation of oxaline. Further structure elucidation via mass spectrometry and nuclear magnetic resonance is required to definitively determine the structures of these analogues.

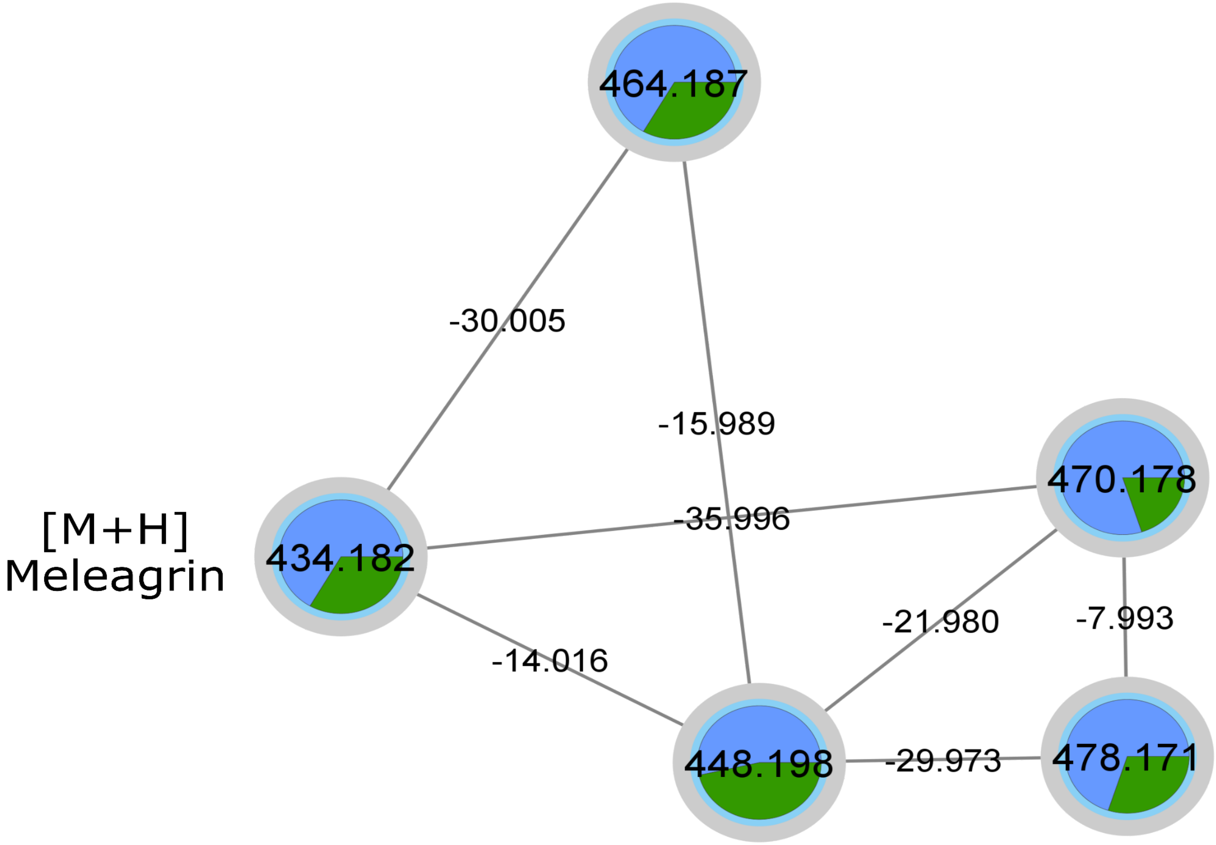


**Figure S1**: Cluster from classical molecular network obtained via GNPS for meleagrin and oxaline adducts and putative analogues. Blue slices represent RS17E extract, while green slices represent RS17F extract.

*Use Case 2: Detection of Secondary Metabolites of Interest in the Presence of Isobars*

*Methods*

Semi-purified fractions RS17E and RS17F generated in *Use Case 1* were analyzed via MALDI-TIMS-qTOF DD. All disposable plastics used were rinsed with methanol prior to contact with fractions, which were then resuspended to a concentration of 1 mg/mL in methanol. Each fraction (1 μL) was co-crystallized with 1 μL of a 1:1 mixture of recrystallized δ-cyano-4-hydroxycinnamic acid (CHCA) and 2,5-dihydroxybenzoic acid (DHB) (Sigma) on an MSP

96-target ground-steel target plate and allowed to air dry. The target plate was then introduced into the MALDI-TIMS-qTOF mass spectrometer and data were acquired via timsControl v2.0.51.0_9669_1571 and flexImaging 5.1 software. The laser power was set to 25% with a frequency of 5000 Hz. The range of detection was from 100 to 2000 Da in positive mode and in TIMS enabled acquisitions, the TIMS range ($1/K_0$) was set to 0.50 to 1.60. Number of laser shots was set to 5000 shots.

*Results and Discussion*

Oftentimes, MALDI DD MS is a relatively quick method to check for the presence of specific metabolites of interest. However, the presence of isobars to metabolites of interest or introduction of isobaric contaminants can often convolute data dependent MS/MS-based structural elucidation efforts. Use of TIMS overcomes these issues by providing additional dimensionality that can be achieved much more quickly than via LC/retention time. Using the same fractions analyzed in *Use Case 1*, **Figure S2** shows a *m/z* 656.062 acquired via MALDI-qTOF MS DD from a fractionated extract that appeared to be differentially oxidated analogues of a secondary metabolite produced by *P. atramentosum* RS17. Upon closer inspection with TIMS enabled, *m/z* 656.062 was found to consist of two different chemical species with $1/K_0$ values of 1.259 and 1.295 (**Figure S2**). With this information, the presence of metabolite(s) of interest can be validated using MALDI-TIMS-qTOF DD and software such as Mass Query Language. Similarly, LC-TIMS-MS/MS in data independent acquisition mode can be used to fragment only the metabolite within a specified *m/z* and $1/K_0$ window.

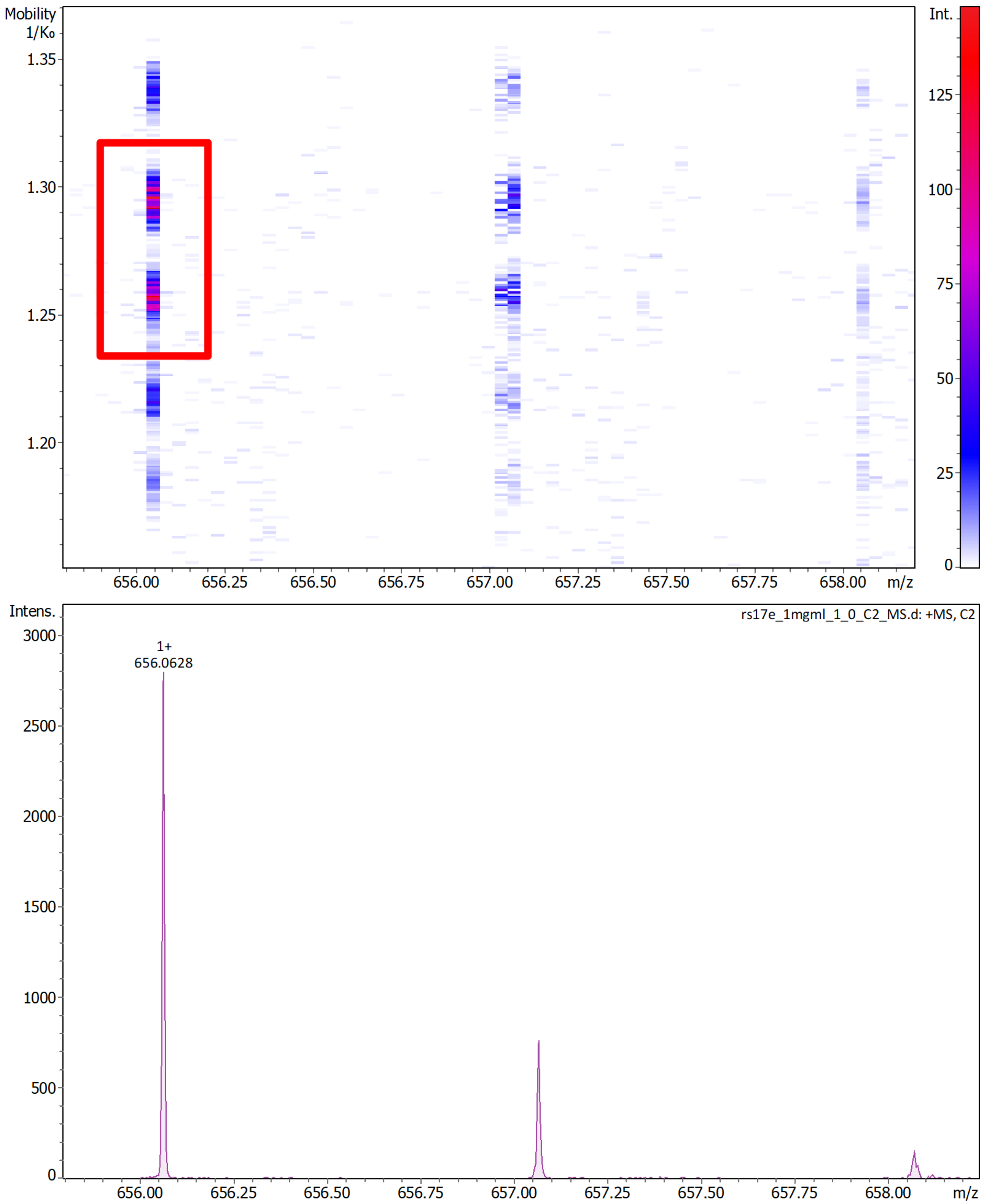


**Figure S2**: Top: Ion mobilogram showing two distinct isobaric species for *m/z* 656.062 at 1.259 and 1.295 $1/K_0$ (highlighted in the red box). Bottom: Zoomed in mass spectrum for *m/z* 656.062 and its isotopologues.

*Use Case 3: Spatial Distribution Secondary Metabolites of Vibrio cholerae rugose Variants using MALDI-qTOF MSI and MALDI-TIMS-qTOF MSI*

*Methods*

*Vibrio cholerae* rugose (O1 El Tor A1552, rugose variant, Rif$^r$) was plated on LB agar and grown overnight at 30 °C (Yildiz and Schoolnik, 1999). A single colony was then used to inoculate a 5 mL LB liquid culture of *V. cholerae* rugose and grown overnight at 30 °C with shaking at 225 rpm. In the MALDI-qTOF MSI run, overnight liquid cultures (5 µL) were spotted on thin agar plates (10 mL of agar in a 90 mm plate) of LB agar media containing 1 mM of taurocholic acid (TCA) or the vehicle (methanol). TCA was purchased from Sigma-Aldrich (≥98%). In the MALDI-TIMS-qTOF MSI run, overnight liquid cultures (5 µL) were spotted on thin agar plates (10 mL of agar in a 90 mm plate) of LB agar media. Biofilm colonies were incubated for 72 h at 30 °C. Following 72 h of growth, colonies were excised from the agar plates using a razor blade and transferred to an MSP 96-target ground-steel target plate (Bruker Daltonics). An optical image of the colonies on the target plate was taken prior to matrix application. A 53 µm stainless steel sieve (Hogentogler Inc.) was used to coat the steel target plate and colonies with MALDI matrix. The MALDI matrix used for the analysis was a 1:1 mixture of recrystallized CHCA:DHB. The plate was then placed in an oven at 37 °C for approximately 4 h or until the agar was fully desiccated. After 4 h, the excess matrix was removed from the target plate and sample with a gentle stream of air. Another optical image was taken of the desiccated colonies on the target plate. The target plate and desiccated colony were then introduced into the MALDI-TIMS-qTOF mass spectrometer and data were acquired via timsControl v2.0.51.0_9669_1571 and flexImaging 5.1 software. The laser power was set to 30%. The range of detection was from 200 to 1500 Da in positive mode. The raster size was set to 1000 μm, and the laser was set to 200 shots per raster point. In the run with TIMS enabled, the $1/K_0$ range was set between 0.78 and 1.60.

*Results and Discussion*

MSI allows for the acquisition of data that provides information on the spatial distribution of metabolites, and metabolites of interest can be identified via manual analysis or automated statistical methods. Here, MSI data for *V. cholerae* rugose variants were acquired via MALDI-TIMS-qTOF MSI and analyzed using a multivariate segmentation workflow in Cardinal MSI previously reported by Luu and Condren *et al.* (Luu et al., 2020). Pixels in this dataset were grouped into one of three segments: inner colony, outer colony/excreted metabolites, and agar control (**Figure S3A**). One of the features, *m/z* 714.2064 was found to be localized to the inner colony, signifying that it may be a bacterial metabolite of interest (**Figure S3B**). The same workflow was used to acquire and analyze data via MALDI-qTOF MSI for *V. cholerae* rugose variants grown in the presence of taurocholic acid (TCA). Interestingly, the chemistry in the inner colony here was found to be similar to the chemistry in the media segments, likely due to decreased bacterial biomass in the center of the biofilm colony (**Figure S3C**). It should be noted that since ion mobility data is not represented in imzML files converted using TIMSCONVERT, converted data are in the same format regardless of whether TIMS is enabled or not during data acquisition. Code used to analyse these data and create figures can be found at https://github.com/gtluu/timsconvert_manuscript_analysis.

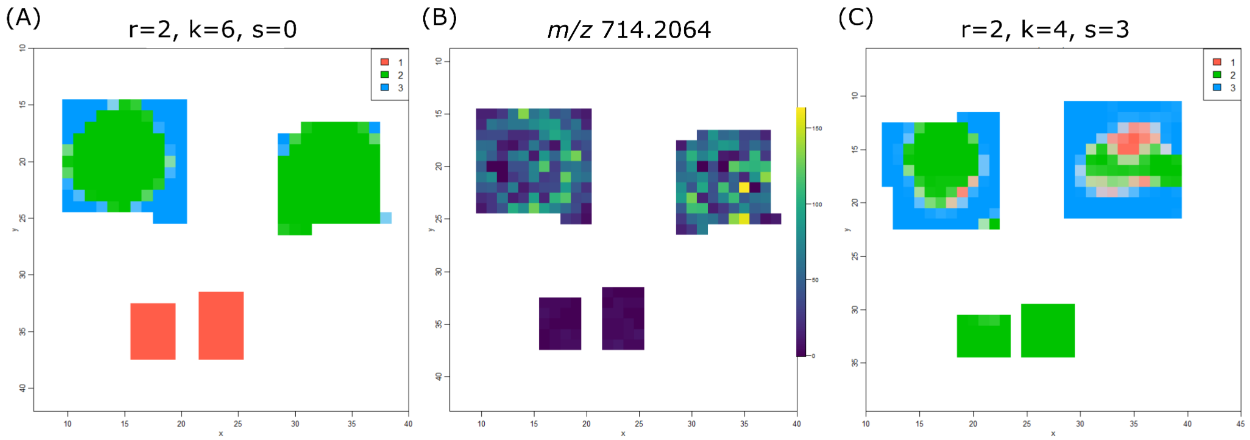


**Figure S3**: (A) Unsupervised segmentation map of untreated *V. cholerae* rugose colonies (MALDI-TIMS-qTOF MSI) generated using Cardinal MSI's spatial shrunken centroids algorithm. (B) Ion image of *m/z* 714.2064 in untreated *V. cholerae* rugose colonies (MALDI-TIMS-qTOF MSI). (C) Unsupervised segmentation map of *V. cholerae* rugose colonies treated with 1 mM TCA (MALDI-qTOF MSI) generated using Cardinal MSI's spatial shrunken centroids algorithm.

### *Data Availability*

Raw datasets analyzed in this manuscript can be found using MassIVE accession MSV000088438.
